# Astrocytic Activation Completely Corrects Memory Performance in an Alzheimer’s Disease Model

**DOI:** 10.1101/2025.09.29.679133

**Authors:** Tirzah Kreisel, Yaara Weinstein, Maya Groysman, Adi Doron, Ron Refaeli, Mickey London, Inbal Goshen

**Author notes:** To whom correspondence should be addressed: Inbal Goshen, Ph.D. Edmond and Lily Safra Center for Brain Sciences (ELSC) The Hebrew University.

## Abstract

Neural and glial dysfunction are thought to underlie memory impairments in Alzheimer’s disease (AD). Conversely, astrocytes are known to play a central role in the healthy brain, and their activation has been shown to enhance memory. To test whether activation of astrocytes can correct memory deficits associated with AD, we expressed hM3Dq in CA1 astrocytes of 7 months old 5XFAD AD model mice, enabling astrocytic chemogenetic Gq pathway activation, following exposure to CNO. We report that astrocytic activation results in complete memory correction, and even memory improvement of AD model mice beyond that of untreated WT levels. We went further to discover the mechanisms underlying this memory improvement in AD mice: First, using 2-photon imaging and electrophysiology, we observed increased neuronal activity (displayed as elevated cFos expression and amplified awake Ca^2+^ events) as well as augmented *in-vivo* long term plasticity following astrocytic activation. Subsequently, we observed increased astrocytic endocytosis of Aβ plaques. Finally, we report that the beneficial effects of astrocytic activation remained stable for an entire year, and persisted beyond the window of astrocytic Gq activation. Our findings that astrocytes correct memory impairments in an AD model could have important clinical implications towards treating this disease.

## INTRODUCTION

Memories define who we are and how we experience the world. Inability to form and access these memories, symptomatic of neurodegenerative conditions such as Alzheimer’s disease (AD), significantly degrades the quality of life. Unfortunately, medical treatments to prevent, hinder, or reverse memory deficits are non-existent. AD causes progressive and irreversible cognitive impairments thought to be consequential of cellular dysfunction. Robust activation and recruitment of non-neuronal cells, including astrocytes, to the areas of neuropathology (Brandebura et al., 2023; Patani et al., 2023; Santello et al., 2019) is a common feature in AD pathology. Indeed, astrocytic involvement in AD was repeatedly hypothesized based on correlative studies showing abnormalities in their number and function in both human patients and animal models of AD (Bouvier et al., 2016; Brandebura et al., 2023; Patani et al., 2023; Santello et al., 2019), together with the fact that many of the physical symptoms accompanying this disease suggest impairments to the classic roles of astrocytes (e.g. monitoring of glutamate levels, neurotrophic factor secretion, and glucose metabolism)(Brandebura et al., 2023; Le Douce et al., 2020; Patani et al., 2023; Santello et al., 2019). Furthermore, reactive astrocytes are found around beta amyloid (Aβ) plaques, and large concentrations of Aβ are found inside astrocytes (Gomez-Arboledas et al., 2018; Nagele et al., 2003; Thal et al., 2000; Zysk et al., 2023), inducing impaired astrocytic function (Allaman et al., 2011; Kuchibhotla et al., 2009; Mei et al., 2010; Wyss-Coray et al., 2003). Such data suggest that alterations in astrocytes modify neural circuit function and underlie AD symptomology (Bouvier and Murai, 2015; Lines et al., 2022; Nanclares et al., 2021). The common conception is that reactive astrocytes contribute to disease progression, and therefore most papers suggest that astrocyte-associated AD treatments should focus on inhibition of astrocyte activity ^e.g.^(Rodriguez-Giraldo et al., 2022).

On the other hand, astrocytes play a central role in normal brain homeostasis in the healthy brain, and pioneering research has shown that these cells can also sense and modify synaptic activity as an integral part of the ’tripartite synapse’ (Araque et al., 1999). The integration of chemogenetic tools (Roth, 2016) in astrocyte research allows real-time, reversible manipulation of intracellular signaling of genetically defined populations in combination with behavior, imaging and electrophysiology. The designer Gq-coupled receptor (hM3Dq) was engineered not to respond to innate ligands, but only to inert drugs like CNO, and can be expressed only in astrocytes under the GFAP promoter. It has a potentiating effect on astrocytic Ca^2+^ events, which serve as a proxy for their activity (Adamsky et al., 2018; Durkee et al., 2019; Van Den Herrewegen et al., 2021), and was shown to induce *de-novo* long term potentiation (LTP) in neurons without any other stimulus (Adamsky et al., 2018; Van Den Herrewegen et al., 2021). Importantly, chemogenetic and optogenetic Gq pathway activation in CA1 hippocampal astrocytes during memory acquisition has been shown to enhance memory in mice (Adamsky et al., 2018; Mederos et al., 2019). Therefore, in this study, we aimed to test whether chemogenetic Gq pathway activation can alleviate AD symptoms.

To explore the possible supportive role of astrocytes in AD, we activated the Gq pathway of dorsal CA1 hippocampal astrocytes in 5XFAD mice (a common AD model) that have five familial AD mutations (Oakley et al., 2006). We chose to test 7 months old mice that already exhibit the full constellation of neurodegenerative symptoms: abnormal hippocampal network dynamics, including unstable memory ensemble traces (Binyamin et al., 2019; Forner et al., 2021; Oakley et al., 2006); reduced LTP (Forest et al., 2021; Forner et al., 2021; MacPherson et al., 2017); and Aβ plaques (Oakley et al., 2006). We examined the effect of astrocyte activation on behavior, as well as its underlying mechanisms. We found that astrocytic activation resulted in a complete correction of memory performance, via increased neuronal activity and plasticity and endocytosis of Aβ plaques. These effects remained for a year, and were long-lasting beyond the period of activation by CNO.

## RESULTS

### Astrocytic chemogenetic activation completely recovers memory in an Alzheimer’s disease model

Previous studies have demonstrated that activation of the Gq-coupled pathway in hippocampal astrocytes during memory acquisition enhances recall in normal mice performing various tasks (Adamsky et al., 2018; Mederos et al., 2019; Refaeli et al., 2024), motivating the question of whether activating the same pathway would improve cognitive function in an Alzheimer’s disease model.

To chemogenetically manipulate the Gq-coupled pathway in CA1 astrocytes, we chose to express the Gq-coupled designer receptor hM3Dq (Roth, 2016), allowing for time-restricted activation by clozapine-N-oxide (CNO). We chose to use 7 months old 5XFAD mice, as they are known to show the full disease phenotype of both cognitive deficiency and Aβ plaque accumulation at this age (Oakley et al., 2006).

We injected 700nl of AAV1 vector encoding hM3Dq fused to mCherry under the control of the astrocytic GFAP promoter (AAV1-gfaABC1D:hM3Dq-mCherry) to the dorsal CA1 (Figure 1A-B). Within the virally-transduced region, hM3Dq expression was limited to the astrocytic outer membranes (Figure S1A), with high penetrance (>98% of the GFAP cells expressed hM3Dq; Figure S1B) and almost complete specificity (>98% hM3Dq positive cells were also GFAP positive; Figure S1C). Co-staining with neuronal nuclei (NeuN) showed no overlap with hM3Dq expression (Figure S1D-E). Similarly, the staining for microglial marker Iba1 revealed no co-localization with hM3Dq (Figure S1F-G). Fear conditioning, in which the manipulation with hM3Dq is most established (Adamsky et al., 2018; Refaeli et al., 2024), could not be performed in 5XFAD mice, as they have a tendency to freeze even before any foot-shock is given, compared to WT (t_(15)_=4.442, p=0.00475)(Figure S1H).

Seven months old 5XFAD and WT littermates were administered CNO via their drinking water for two weeks, (3mg/kg/day) while controls were given regular water. They were then all trained in the radial arm water maze (RAWM) for 4 days and tested on the fifth day (Figure 1C), where every entrance to a wrong arm was counted as an error. Seven months old 5XFAD mice that received regular water, performed worse than WT controls (F_(1,40)_=16.512, p=0.000220)(Figure 1D), as was previously shown in this and other tasks (Binyamin et al., 2019; O’Leary and Brown, 2022; Priori et al., 2023). After 2 weeks of CNO administration, memory was enhanced in WT mice expressing hM3Dq in CA1 astrocytes (p=0.05), as we have shown in other tasks before (Adamsky et al., 2018; Refaeli et al., 2024). Remarkably, CNO activation of hM3Dq in CA1 astrocytes of 5XFAD mice enhanced memory (p=3.8E^-5^) to the same level, such that no difference could be observed between CNO-receiving 5XFAD and WT groups (p=0.565) (Figure 1D).

**Figure 1:**
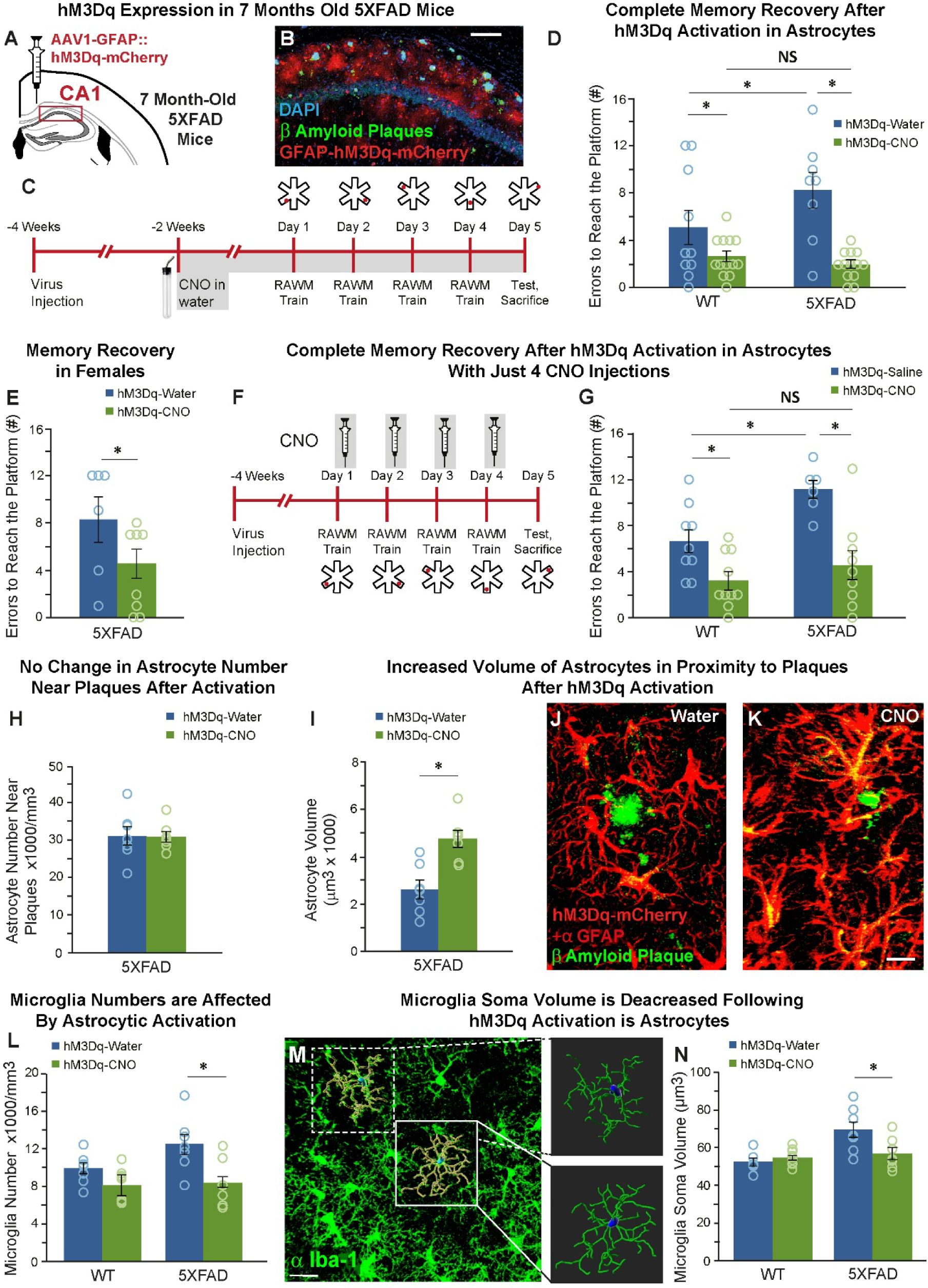
Complete memory recovery in 5XFAD mice following astrocytic chemogenetic activation. (**A**) Bilateral double injection of AAV1-GFAP::hM3Dq-mCherry to the CA1 of 7 month old 5XFAD or WT mice, (**B**) resulted in hM3Dq expression (red) selectively in CA1 astrocytic membrane around the soma, as well as in the distal processes CA1 astrocytes. Aβ plaque are visible (green). Scale bar = 100µm. (**C**) The experiment schedule. (**D**) Mice expressing hM3Dq in their CA1 astrocytes drank either water (WT n=10; 5XFAD n=8) or CNO (WT n=13; 5XFAD n=13). 5XFAD-water showed 60% more errors than WT-water (*= p< 0.00025). Astrocytic activation by CNO application resulted in more than 40% improvement in the number of errors, compared to water treated WT (*= p<0.05), and more than 70% improvement in water treated 5XFAD (*= p<0.00005). (**E**) Astrocytic activation by CNO application resulted in more than 40% improvement in the number of errors in 5XFAD female mice (n=8), compared to water treated 5XFAD female WTs (n=6) (*= p<0.05). (**F**) The experiment schedule. (**G**) Mice expressing hM3Dq in their CA1 astrocytes were injected with either saline (WT n=9; 5XFAD n=6) or CNO (WT n=10; 5XFAD n=9). 5XFAD-saline showed 65% more errors than WT-saline (*= p< 0.01). Astrocytic activation by CNO application resulted in more than 50% improvement in the number of errors, compared to saline injected WT (*= p<0.02), and more than 60% improvement in water treated 5XFAD (*= p<0.00025). (**H**) There was no effect on the number of astrocytes proximal to Aβ plaque in hM3Dq-expressing 5XFAD mice that drank either water (n=7) or CNO (n=7) for 2 weeks. (**I-K**) Astrocytes volume in proximity to Aβ plaque was increased in hM3Dq-expressing 5XFAD mice that drank CNO (n=7) compared to the those who drank Water (n=7) (*= p<0.000845). Staining for GFAP (red) and Aβ plaques (green). Scale bar = 10µm. (**L**) CNO application decreased microglia number in 5XFAD mice (Water n=7; CNO n=6) (*= p< 0.02), but had no effect on WT mice (Water n=7; CNO n=9). (**M**) Microglia were stained by an α-Iba-1 antibody (green; left). A 3D stack was analyzed by the IMARIS program (right). Scale bar = 20µm. (**N**) CNO application had no effect on WT mice (Water n=6; CNO n=10), but decreased microglia soma volume in 5XFAD mice (Water n=7; CNO n=7) (*= p< 0.025). Data presented as mean ± standard error of the mean (SEM).

To verify that astrocytic activation does not have a direct effect on swimming speed, we tested it in the same GFAP::hM3Dq mice, after the first day of RAWM training. Importantly, no difference in velocity was found between the groups (F_(1,41)_=1.085, p=0.304)(Figure S1I). To confirm that our results did not stem from the CNO application itself, we injected an additional cohort of WT and 5XFAD mice with a control AAV1-gfaABC1D::tdTomato vector. CNO application in these control mice had no effect on performance in the RAWM (Figure S1J), with 5XFAD mice showing, as expected, an increased number of errors (F_(1,16)_=10.111, p=0.006).

Given the higher prevalence of AD in women (2021; Cao et al., 2020), and the behavioral differences found between males and females in 5XFAD mice (Forner et al., 2021), it was important to repeat our experiment in females. We found that activation of hM3Dq in CA1 astrocytes of female 5XFAD mice improved their memory in the RAWM (t_(12)_=1.959, p=0.037)(Figure 1E).

We then tested whether an even shorter-term CNO application would have a similar effect to 2 weeks of CNO drinking. 7 months old 5XFAD and WT littermates were administered CNO via IP injections (3mg/kg, controls were given saline) for only 4 days, during which RAWM was acquired, and then tested on the 5^th^ day (Figure 1F). In the 7 months old mice that received saline, the 5XFAD mice performed worse than WT (F_(1,30)_=23.026, p=0.001)(Figure 1G). After 4 days of CNO injections, hM3Dq activation in CA1 astrocytes enhanced memory in WT mice (p=0.018). Remarkably, hM3Dq activation in CA1 astrocytes of 5XFAD mice enhanced memory (p=2.35E^-4^) to the same level, such that no difference was observed between CNO-receiving 5XFAD and WT groups (p=0.334) (Figure 1G), just as we have shown after two weeks of CNO administration. Swimming velocity was monitored to ensure that astrocytic activation did not have a direct effect on swimming speed, and, again, no difference in velocity was found between the groups (F_(1,30)_=1.413, p=0.244)(Figure S1K).

We proceeded to check whether the astrocytes or microglia cells in 5XFAD mice differ in number or morphology from those of their WT littermates, and whether hM3Dq activation can affect these changes. After 2 weeks of hM3Dq activation, no differences were found in the overall astrocyte number (F_(1,27)_=0.446, p=0.51)(Figure S1L), soma volume (F_(1,27)_=0.002, p=0.964)(Figure S1M) or processes length (F_(1,27)_=0.049, p=0.826)(Figure S1N). Several papers reported changes in astrocytes proximal to the plaques in other models of AD ^e.g.^(Bouvier et al., 2016; Reichenbach et al., 2019). In our experiment, (50um) astrocyte number remained the same in proximity to the plaques (t(12)=0.0778, p=0.469)(Figure 1H), however astrocytic volume was significantly increased (t(12)=4.023, p=0.000845)(Figure 1I-K). It should be noted that Figures 1J and 1K are in the same magnification. Additionally, 2 weeks of astrocyte hM3Dq activation lowered microglia cell numbers (F_(1,25)_=11.893, p=0.002), especially in 5XFAD mice (p=0.017)(Figure 1L). Soma volume was initially increased in 5XFAD mice (F_(1,26)_=11.093, p=0.003) and decreased after astrocytic manipulation (p=0.025)(Figure 1M-N). No changes were found in microglia processes length (F_(1,26)_=0.279, p=0.602)(Figure S1O).

Our results show that astrocytic activation affects morphological traits of astrocytes and microglia in an Alzheimer’s disease model. They further demonstrate that astrocytic Gq-coupled pathway activation in two different protocols, lead to a complete recovery of memory performance. Unexpectedly, the memory of treated 5XFAD mice became as good as that of the treated controls.

### Chemogenetic Gq-pathway activation in astrocytes enhances neuronal activity and plasticity in an Alzheimer’s disease model

Astrocytes can influence memory via their effect on neurons, and neuronal dysfunction is one of the hallmark symptom of AD (Palop and Mucke, 2016). Specifically, Gq pathway activation in astrocytes has an activity-dependent effect on neuronal activity, i.e. the increase in neuronal activity occurs only when astrocytic activation is paired with learning (Adamsky et al., 2018). Recently, it was shown that the immediate early gene (IEG) Arc is reduced following fear conditioning in the Alzheimer’s disease model APP/PS1(Perusini et al., 2017). We used another IEG, cFos, as a quantifiable proxy to neuronal activity, because it is elevated when a neuron is hyper-activated during a memory task. We found that astrocytic activation via hM3Dq enhanced cFos levels 90 min after the last RAWM memory test (F_(1,26)_=13.092, p=0.001), especially in 5XFAD mice (p=0.01)(Figure 2A-B). Astrocytic activation via hM3Dq enhanced cFos levels 90 min after the last RAWM memory test in females as well (t_(9)_= 2.178, p=0.0286)(Figure S2A), and 4 days of CNO injection causing astrocytic activation via hM3Dq enhanced neuronal cFos levels 90 min after the last RAWM memory test (F_(1,29)_=4.786, p=0.037)(Figure S2B).

cFos is a one time-point proxy to neuronal activity, and based on these results we wanted to examine continuous neuronal activity in real-time. Earlier studies showed an altered spontaneous neuronal activity *in-vivo* in the CA1 of 5XFAD mice. Specifically, they demonstrated hyper-excitability in awake (Yao et al., 2020) and anesthetised or sleeping mice (Zarhin et al., 2022). We expressed AAV5-CaMKII::GCaMP6f and imaged spontaneous activity in CA1 pyramidal neurons using a 2-Photon microscope (Figure 2C). In our hands, 7 months old anesthetised 5XFAD mice showed an increased number of events compared to WT controls (t_(10)_=2.126, p=0.03)(Figure S2C-D).

**Figure 2:**
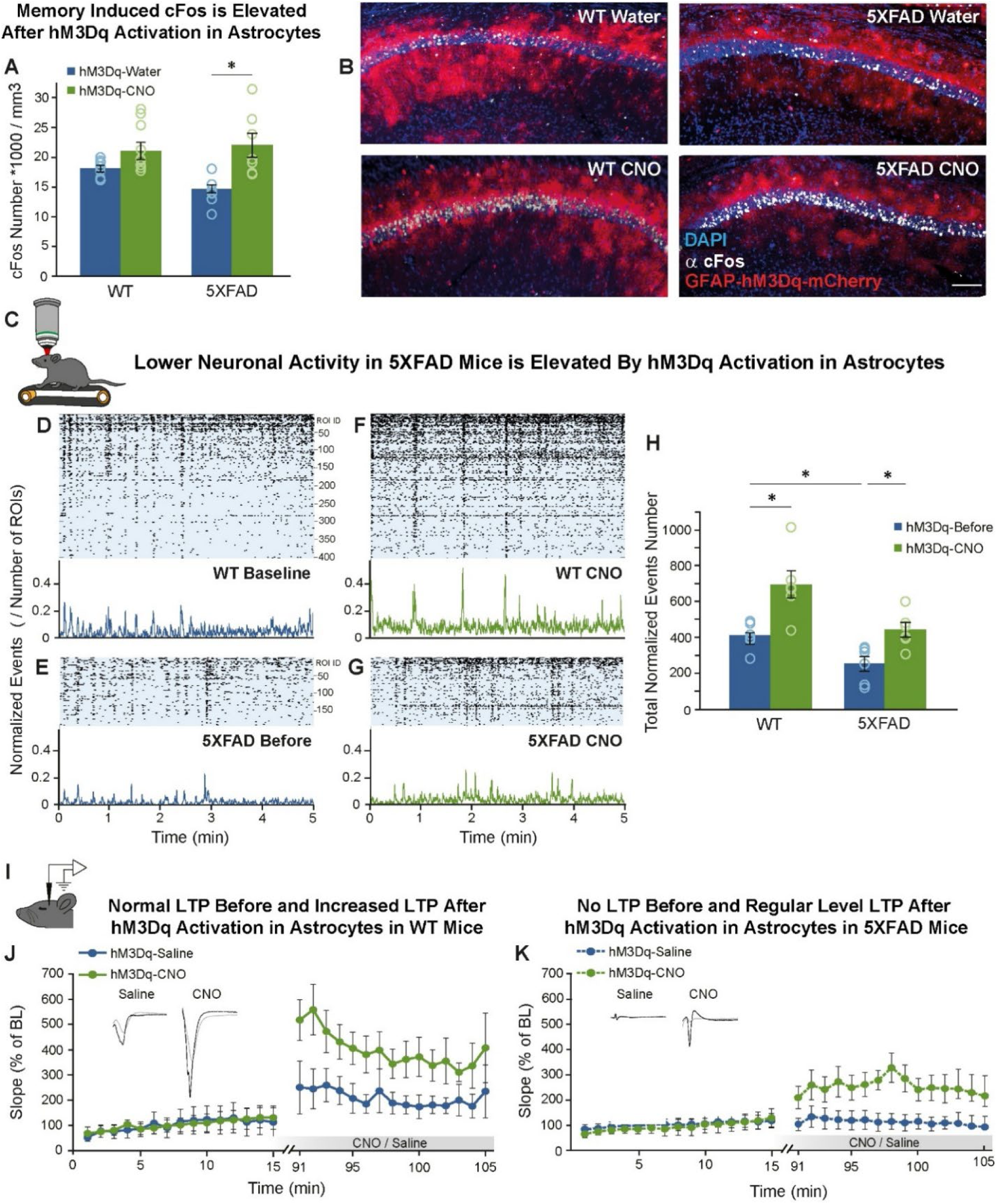
Neuronal activity and plasticity are enhanced in 5XFAD mice following astrocytic chemogenetic activation. (**A**) CNO application elevated cFos levels overall (p=0.001). Post hoc analysis showed that CNO significantly elevated cFos levels in 5XFAD mice (Water n=7; CNO n=7) (*= p< 0.01), but not in WT mice (Water n=7; CNO n=9). (**B**) Representative images from all groups, showing hM3Dq-mCerry (red), cFos (white) and nuclei (blue). Scale bar 100µm. (**C**) Neuronal activity was measured as Ca^2+^ events, *in-vivo* in awake mice. Representative traces of normalized neuronal activity (events/ROIs number) below, of the ROIs above, in 5 min of WT (**D**) and 5XFAD (**E**) before, and WT (**F**) and 5XFAD (**G**) after CNO. (**H**) The neuronal activity in mice expressing hM3Dq in their CA1 astrocytes (WT n=6; 5XFAD n=6) was measured before and after CNO injection. 5XFAD-Before showed over 35% less activity than WT-Before (*= p< 0.04). Astrocytic activation by CNO application resulted in more than 65% elevation in the number of event in WT mice (*= p<0.00075), and more than 70% elevation in 5XFAD (*= p<0.015). (**I**) LTP measured *in-vivo* was induced by HFS. (**J**) HFS induced LTP (>200% increase) in saline injected WT (n=7; left insert: before HFS in gray, after HFS in black). The same HFS induced a larger LTP (>400% increase) in CNO injected WT (n=7; right insert). (**K**) HFS did not induce LTP in saline injected 5XFAD (n=5; left insert: before in gray, after in black). The same HFS induced LTP (>250% increase) in CNO injected 5XFAD (n=8; right insert). Data presented as mean ± standard error of the mean (SEM).

We then expressed AAV5-CaMKII::GCaMP6f in CA1 pyramidal neurons and AAV1-gfaABC1D::hM3Dq-mCherry in CA1 astrocytes (Figure S2E), and imaged spontaneous neuronal activity before and after CNO application. When activity in awake mice was imaged, a different picture emerged: CNO injection increased neuronal activation in both WT and 5XFAD mice (F_(1,10)_=34.980, p=0.000148). Prior to CNO application, 5XFAD mice showed comparatively reduced numbers of neuronal Ca^2+^ events (p=0.04) (Figure 2D-E,H). CNO application augmented the number of events in both WT (p=0.000745)(Figure 2F,H) and 5XFAD mice (p=0.015)(Figure 2G,H), and furthermore increased synchronous activity (the number of concurrent events) in both groups (F_(1,10)_=17.3666, p=0.002)(Figure S2F-G, 2D-G).

Finally, after finding evidence of changes in neuronal activity, we proceeded to explore neuronal plasticity. Previous studies have shown decreased LTP in 5XFAD mice (Forest et al., 2021; Forner et al., 2021; MacPherson et al., 2017). We and others have shown in the past that astrocytic Gq-pathway activation can induce neuronal *de-novo* LTP in brain slices (Adamsky et al., 2018; Mederos et al., 2019; Van Den Herrewegen et al., 2021), and OptoGq can increase LTP in slices (Gerasimov et al., 2023). Given these results, we examined LTP induction after hM3Dq activation by CNO injection, in 7-month-old 5XFAD and WT mice. Specifically, we performed *in-vivo* recordings before and after delivering high frequency stimulation (HFS; five trains of 200Hz for 100ms, presented six times at 1min intervals) to generate LTP (Figure 2I). CNO (3mg/kg) was given after baseline measurement, 30 minutes before the HFS. Following HFS, saline injected WT controls expressed normal LTP (>200% increase), and CNO treated mice showed a higher LTP (>400% increase), which resulted in a significant difference between groups (F_(29,232)_=2.764, p=1.27E^-5^)(Figure 2J). In 5XFAD mice, HFS did not induce LTP in the saline injected group (Figure 2K, blue), as was previously shown (Forest et al., 2021; Forner et al., 2021; MacPherson et al., 2017). CNO administration in 5XFAD mice caused an expression of LTP (>250% increase), significantly different from saline (F_(29,145)_=10.721, p=1.06E^-23^)(Figure 2K, green).

Our results show that activation of the Gq-coupled pathway in astrocytes promotes neuronal activity and stimulates neuronal plasticity in 5XFAD Alzheimer’s model mice, which can explain the marked memory recovery in these mice.

### Chemogenetic Gq-pathway activation increases astrocytic endocytosis of Aβ plaques in an Alzheimer’s disease model

Wo took an additional path in order to explain our strong behavioral results, and examined whether recruitment of the Gq-coupled pathway in astrocytes influenced the main physiological marker of AD, Aβ plaques. Remarkably, we found that 2 weeks of hM3Dq activation caused a 40% reduction in Aβ plaque volume in the CA1 of 5XFAD mice (t_(11)_=2.9, p=0.014)(Figure 3A-C), but had no effect on plaques in the nearby hippocampal region, the dentate gyrus, where hM3Dq is not expressed (t_(11)_=0.745, p=0.471)(Figure S3A). Control mice are not shown because they have no Aβ plaques. In female 5XFAD mice, Aβ plaque volume was reduced in the CA1 following 2 weeks of hM3Dq activation (t_(14)_=1.921, p=0.0376)(Figure 3D). To verify that CNO application itself does not produce a similar effect, we injected 5XFAD mice with a control virus (AAV1-gfaABC1D:: tdTomato), and found that CNO alone did not affect Aβ plaque volume (Figure S3B).

Upon measuring the Aβ plaques in the 5XFAD mice that showed improved memory after only 4 days of CNO injections (Figure 1F-G), we surprisingly found that plaque volume in the CA1 was already reduced by 50% (t_(12)_=4.156, p=0.001)(Figure 3E). No effect on Aβ plaques was found in the nearby hippocampal region, dentate gyrus, where hM3Dq is not expressed (t_(12)_=1.286, p=0.111) (Figure S3C).

In order to ascertain the cause of the rapid reduction in Aβ plaque volume, we measured the overlap of astrocytes, Aβ plaques, and lysosomes in the CA1 of 5XFAD mice one day after a single injection of saline or CNO (Figure 3F-I). We found that astrocytic activation of hM3Dq with CNO significantly increased endocytosis by astrocytes, as demonstrated by the overlap of lysosomes and Aβ inside astrocytes (t_(9)_=3.787, p=0.002)(Figure 3J). To determine whether endocytosis is also increased in microglia upon astrocytic activation, we tested the overlap of Aβ plaques with microglial lysosomes, but found no difference between CNO and saline injected controls (t_(9)_=0.335, p=0.670)(Figure S3D-E).

The degree of endocytosis can also be seen by the level of Aβ plaque sphericity, as Aβ plaques are naturally spherical, and lose some of their sphericity when endocytosed by cells. Sphericity was found to be decreased in CNO-treated 5XFAD mice compared to saline-treated controls (t_(9)_=3.665, p=0.005)(Figure 3K-M), and, as expected, no significant difference in Aβ plaque volume was found between CNO and saline controls one day after a single injection (t_(9)_=0.398, p=0.700)(Figure S3F).

Finally, these results prompted us to re-evaluate the findings of 2-week CNO administration, because we noticed that the astrocytes around plaques seemed to have Aβ inside them (Figure 1K). Upon further examination of these slides, we indeed found that astrocytic activation of hM3Dq with CNO significantly increases endocytosis by astrocytes, as demonstrated by the overlap of Aβ and GFAP (t_(10)_=3.71, p=0.002)(Figure S3G-I).

Our results show that endocytosis by astrocytes, stimulated by their Gq-coupled pathway, starts just one day after activation in 5XFAD mice.

**Figure 3:**
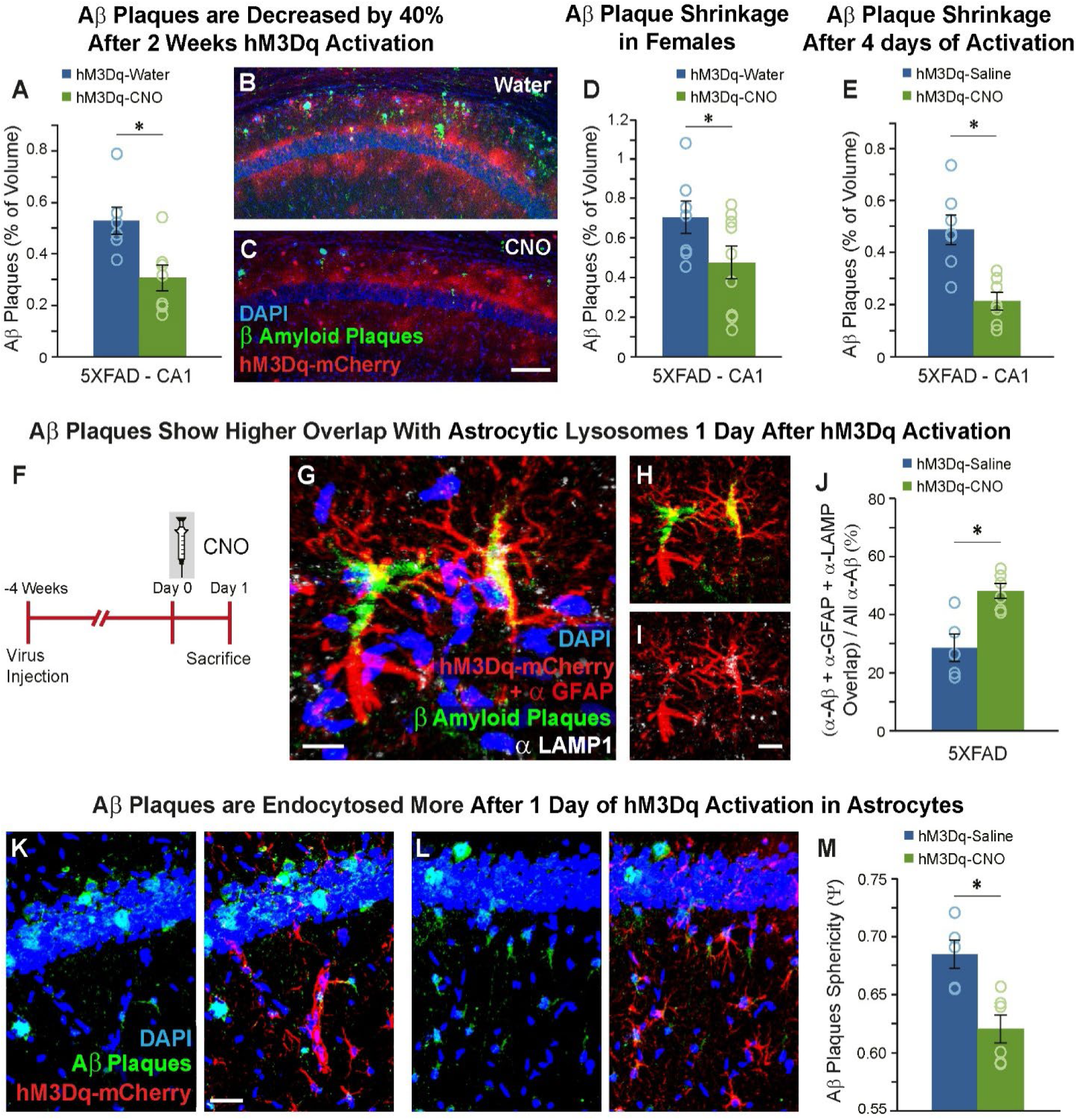
Astrocytic endocytosis of Aβ plaques is induced by astrocytic Gq-pathway activation in an Alzheimer’s disease model. (**A-C**) Two weeks CNO-treated 5XFAD mice (n=7) expressing hM3Dq in CA1 astrocytes showed 40% less Aβ plaque in their CA1 compared to 5XFAD mice that received water (n=6) (*= p< 0.0014). Scale bar = 100µm. (**D**) CNO-treated 5XFAD female mice (n=9) showed 30% less Aβ plaque compared to 5XFAD female mice that received water (n=7) (*= p<0.05). (**E**) Four days CNO-injected 5XFAD mice (n=7) expressing hM3Dq in CA1 astrocytes showed 40% less Aβ plaque in their CA1 compared to 5XFAD mice that were injected with saline (n=7) (*= p< 0.001). (**F**) The experiment schedule. (**G-I**) Expression of hM3Dq + GFAP staining (red), staining for Aβ plaques (green) and lysosomes (α-LAMP1; white). Nuclei are in blue. Scale bar 10µm. (**J**) One day following injection, 5XFAD-CNO (n=5) showed 65% more overlap of astrocytes, Aβ plaques and lysosomes compared to 5XFAD-saline (n=6)(*= p< 0.0025). (**K-L**) Expression of hM3Dq (red) and staining for Aβ plaques (green) in WT (**K**) and 5XFAD (**L**). Nuclei are in blue. Scale bar 30µm. (**M**) One day following injection, 5XFAD-CNO (n=6) showed 65% less Aβ plaques sphericity compared to 5XFAD-saline (n=5)(*= p< 0.005). Data presented as mean ± standard error of the mean (SEM).

### Clinically relevant schedules of Gq-pathway activation recover memory, elevate neuronal activity, and reduce Aβ plaques in an Alzheimer’s disease model

We tested the ability of hM3Dq activation in astrocytes to alleviate AD symptoms in 7 months old mice, in which a complete disease phenotype was displayed, as a model of diagnosed human patients. Our results are promising as they show the ability of astrocyte Gq-pathway activation to improve both the cognitive and physiological deficiencies in this AD model after they have already occurred. However, the short treatment regiment described above is not a realistic one. First, because a lifelong treatment starting at diagnosis is clinically required, and second, since the treated astrocytes may be damaged and even die as a result of prolonged activation.

**Figure 4:**
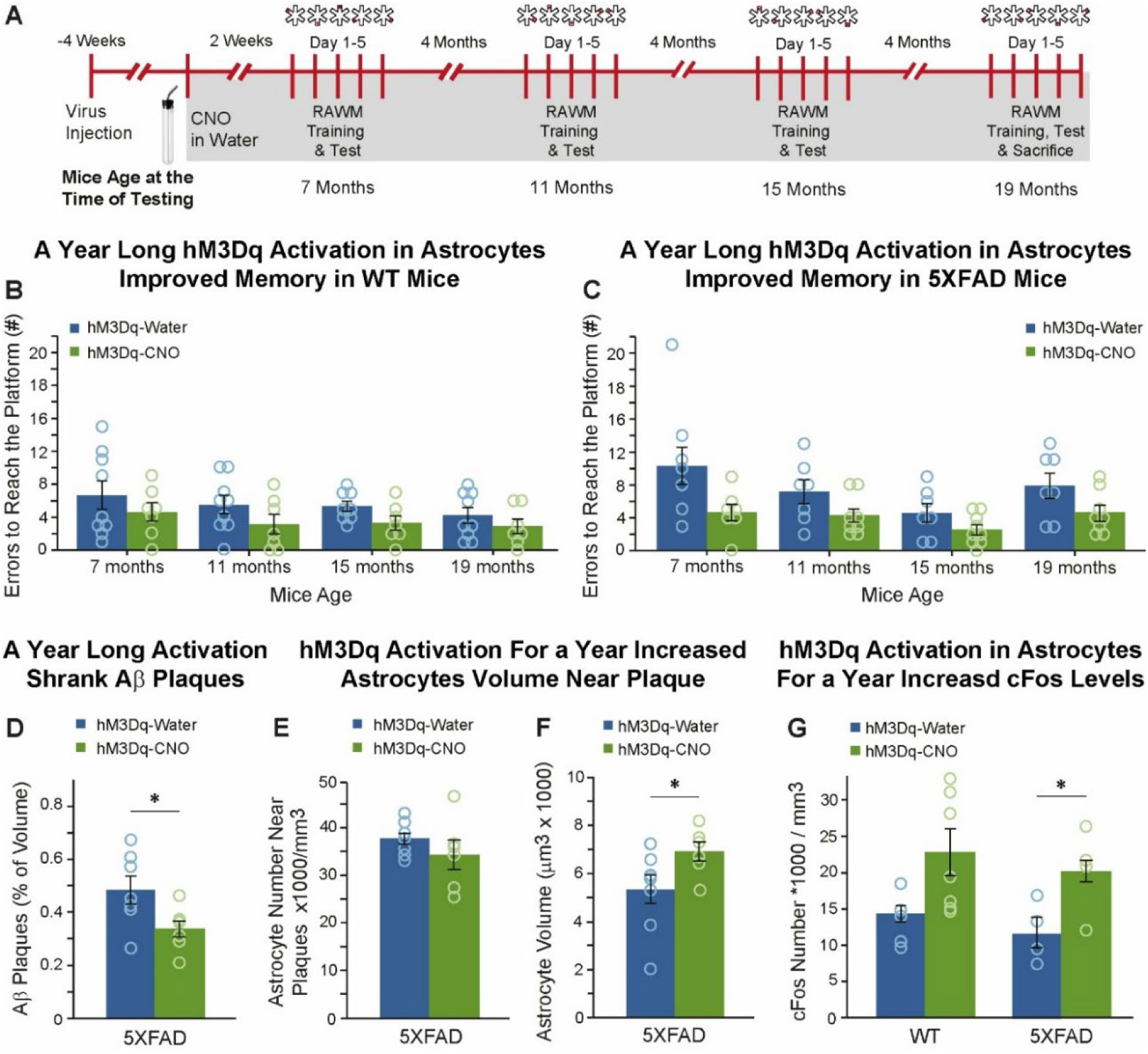
Gq-pathway activation for a full year recovers memory and improves physiological symptoms in an Alzheimer’s disease model. (**A**) The schedule of the year-long experiment. (**B**) During a year of treatment, CNO given WT (n=7) showed less errors than water given WT (n=9)(*= p< 0.05). (**B**) During a year of treatment, 5XFAD given CNO (n=9) showed less errors than 5XFAD given water (n=7)(*= p< 0.01). (**D**) CNO-treated 5XFAD mice (n=7) expressing hM3Dq in CA1 astrocytes showed 30% less Aβ plaque in their CA1 compared to 5XFAD mice that received water (n=7)(*= p< 0.00025). (**E**) There was no effect on the number of astrocytes proximal to Aβ plaque in hM3Dq-expressing 5XFAD mice that drank either Water (n=8) or CNO (n=6) for a year. (**F**) Astrocyte volume in proximity to Aβ plaque was increased in hM3Dq-expressing 5XFAD mice that drank CNO (n=6) compared to the those who drank water (n=8)(*= p<0.031) for a year. Scale bar = 10µm. (**G**) CNO application for a year elevated cFos levels overall (p=0.008). Post hoc analysis showed that CNO significantly elevated cFos levels in 5XFAD mice (Water n=7; CNO n=6)(*= p< 0.05), but not in WT mice (Water n=7; CNO n=6)(p=0.055). Data presented as mean ± standard error of the mean (SEM).

To extend our results and test whether Gq-pathway activation can alleviate AD symptoms for a longer time period, we performed a year-long experiment (Figure 4A): we started, as before, at age 7 months, CNO was continuously given in the drinking water, and RAWM performance was tested every 4 months for a year (i.e., at ages 7, 11, 15, and 19 months). At the end of the year (in 19 months old mice), 90 min after the last memory test, brains were collected and we measured cFos levels and Aβ plaques. Continuous CNO administration for the span of a year, activating hM3Dq in CA1 astrocytes, improved memory in both WT (F_(3, 42)_=3.277, p=0.03)(Figure 4B), and 5XFAD mice (F_(3, 42)_=4.4, p=0.009)(Figure 4C). To verify that prolonged astrocytic activation does not have a direct effect on swimming speed, we tested it at the end of the first day of RAWM in 19 months old mice. No difference in velocity was found between the groups (F_(1,26)_=0.048, p=0.829)(Figure S4A).

In addition, we found that a year of hM3Dq activation caused a 30% reduction in Aβ plaque volume in the CA1 of 5XFAD (t_(12)_=2.419, p=0.016)(Figure 4D). The number of astrocytes remained the same in proximity to the plaques (t(12)=1.11, p=0.144)(Figure 4E), but astrocytic volume in the same areas was increased (t(12)=2.052, p=0.031)(Figure 4F). Finally, we show that year-long astrocytic activation with hM3Dq increased cFos levels, 90 min after the last RAWM memory test at age 19 months (F_(1,22)_=8.640, p=0.008)(Figure 4G).

Another clinically relevant question arises in regards to the long-term effects of Gq-pathway activation if the CNO treatment is only given intermittently, i.e. would the favorable symptoms immediately disappear upon cessation of treatment, or would they linger for some time? To that end, we performed the following experiment (Figure 5A): we administered CNO in the drinking water of 7 months old mice for two weeks, as before, and tested RAWM performance, after which we ceased CNO administration. 2 weeks later, we performed behavioural testing, and some mice were randomly selected for histology to quantify Aβ plaques and cFos. The rest of the mice remained for 4 more weeks (total of 6 weeks with no CNO) and subsequently underwent behavioral testing and histology (Figure 5A).

As we showed earlier, following a 2-week course of CNO, hM3Dq in CA1 astrocytes enhanced memory in WT and in 5XFAD mice (F_(1,48)_=4.85, p=0.032)(Figure 5B) to the same level, such that no difference could be observed between CNO-receiving 5XFAD and WT groups (p=0.619)(Figure 5B). Remarkably, after two weeks *without* CNO, the behavioral result in the same mice remained improved, i.e. former activation of hM3Dq in CA1 astrocytes two weeks earlier still enhanced memory in WT and 5XFAD mice (F_(1,48)_=5.847, p=0.019)(Figure 5B), and no difference was observed between the former CNO-receiving 5XFAD and WT groups (p=0.814)(Figure 5B). Histological measures revealed that the influence on neuronal activity also remained after two weeks without CNO administration, as we found that former astrocytic activation with hM3Dq enhanced cFos levels in WT and 5XFAD mice (F_(1,17)_=10.703, p=0.004)(Figure 5D). In addition, Aβ plaque volume in the CA1 of 5XFAD mice was reduced by 30%, even after two weeks without CNO administration (t_(5)_=2.302, p=0.034)(Figure 5E). However, when the remaining mice were retested after another 4 weeks (6 weeks total) without CNO, the effects of Gq activation had vanished (F_(1,22)_=0.217, p=0.646), so that only the strain had an effect on behavior, with 5XFAD mice in general performing worse (F_(1,22)_=4.167, p=0.05) (Figure 5F). At this stage, no effect of the former CNO treatment on cFos levels could be detected (F_(1,22)_=0.14, p=0.905), and only the strain had an effect, with 5XFAD mice in general showing lower cFos levels (F_(1,22)_=7.137, p=0.014)(Figure 5G). No effect of the former CNO treatment was found on Aβ plaque volume (t_(10)_=4.697, p=0.134)(Figure 5I). Finally, no effect on swimming velocity was found between the groups after 2 weeks with CNO (F_(1,48)_=0.902, p=0.347)(Figure S4B), 2 weeks without CNO (F_(1,48)_=0.002, p=0.965)(Figure S4C), and 6 weeks without CNO (F_(1,22)_=0.088, p=0.770)(Figure S4D).

**Figure 5:**
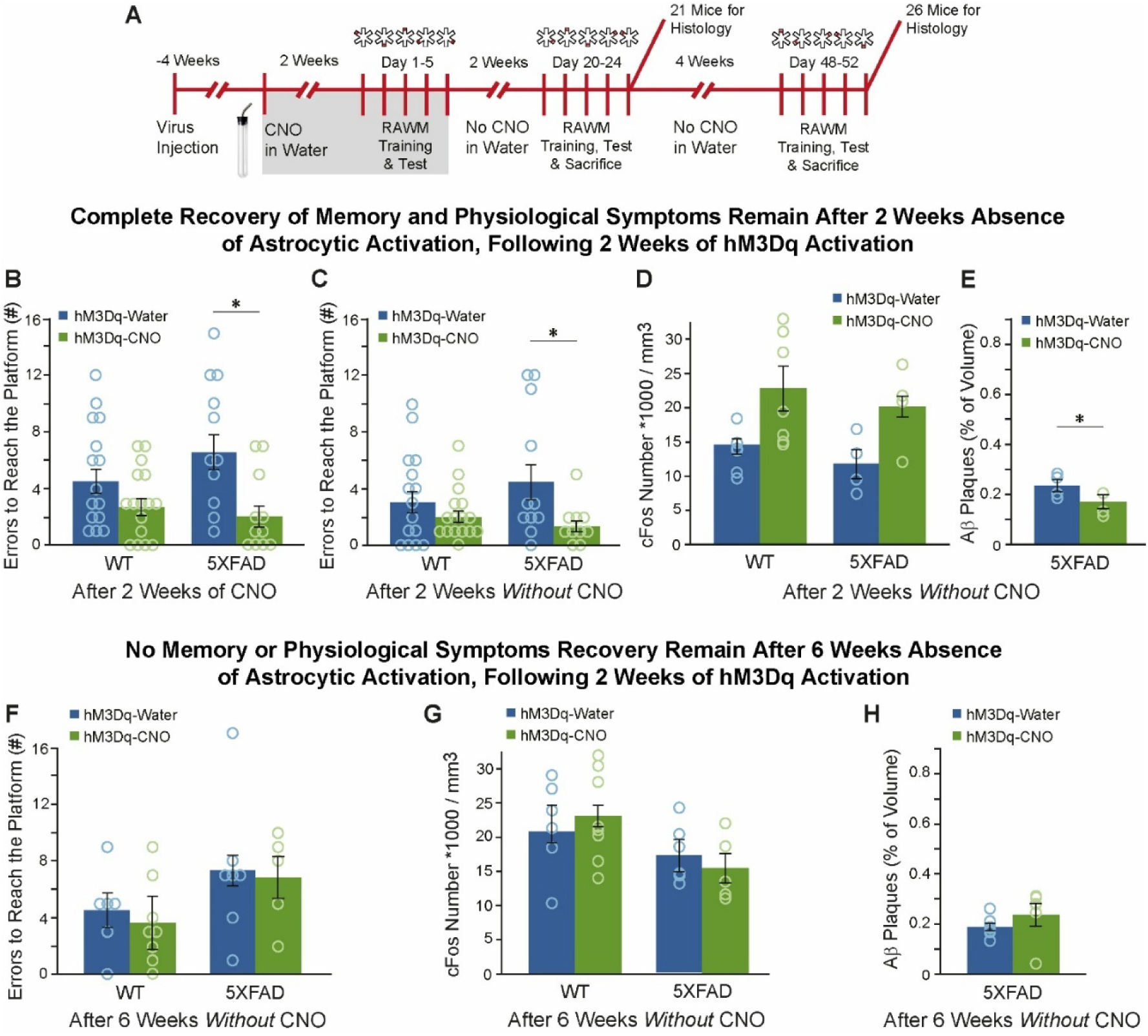
Gq-pathway activation bears long term effects, and can recover memory and improve physiological symptoms in an Alzheimer’s disease model, two weeks after CNO is no longer given. (**A**) The schedule ON-OFF CNO experiment. (**B**) Mice drank either water (WT n=15; 5XFAD n=11) or CNO (WT n=15; 5XFAD n=11) for 2 weeks. Astrocytic activation by CNO application had an effect over all (p=0.038), and post hoc tests showed the more than 65% improvement in water treated 5XFAD is significant (*= p< 0.001). (**C**) After two weeks *without* CNO, the same mice, still showed an effect over all (p=0.019), and post hoc tests showed the 70% improvement in water treated 5XFAD is significant (*= p< 0.01). (**D**) 21 mice were taken for histology at this time point, and it was found that CNO application elevated cFos levels overall (p=0.004). Post hoc analysis showed that CNO significantly elevated cFos levels in WT mice (Water n=6; CNO n=7) (*= p< 0.015), but not in 5XFAD mice (Water n=4; CNO n=4) (*= p< 0.058). (**E**) 5XFAD mice (n=3) expressing hM3Dq in CA1 astrocytes that were treated with CNO for 2 weeks, and spent 2 weeks *without* CNO still showed 25% less Aβ plaque in their CA1 compared to 5XFAD mice that received water (n=4) (*= p< 0.035). (**F**) After four more weeks (total of 6 weeks) *without* CNO, the remaining 26 mice (WT-Water n=6; WT-CNO n=8; 5XFAD-Water n=7; 5XFAD-CNO n=5) did not show any effect on memory performance. (**G**) The same mice showed an overall strain effect (p=0.014), but no effect of the CNO was detected. (**H**) 5XFAD mice expressing hM3Dq in CA1 astrocytes that were treated with CNO for 2 weeks and spent 6 weeks *without* CNO did not show any change in Aβ plaque volume compared to 5XFAD mice that received water. Data presented as mean ± standard error of the mean (SEM).

Our results show that a year-long activation of Gq-coupled pathway in astrocytes improves memory and reduces Aβ plaque burden continually throughout the treatment period. Additionally, we found that when CNO is given intermittently, the positive cognitive and histological effects last for two weeks of CNO absence in 5XFAD Alzheimer’s model mice.

## DISCUSSION

In this work, we demonstrated that activation of the Gq-coupled pathway in astrocytes rectifies many of the neurodegenerative symptoms of AD model mice. Specifically, astrocytic activation completely corrected memory functioning, possibly by augmenting neuronal activity and plasticity, as well as initiating endocytosis of Aβ plaques by astrocytes. Furthermore, we showed that the beneficial effects can persist for a year and are long-lasting, i.e. they continue for a limited time even without astrocytic activation.

Past research in Alzheimer’s disease models in general (Brandebura et al., 2023; Patani et al., 2023; Santello et al., 2019) and 5XFAD mice specifically (Lu et al., 2023; Samokhina et al., 2025; Weiss et al., 2025), has demonstrated impaired astrocytic function. For that reason, studies have focused mainly on blocking astrocyte activity in various ways, assuming that reactive astrocytes are responsible for some of the symptoms ^e.g.^ (Chen et al., 2025; Chicote-Gonzalez et al., 2025; Lee et al., 2023; Nakano-Kobayashi et al., 2023; Reichenbach et al., 2018; Zeng et al., 2023; Zhang et al., 2023), and reported improved cognitive performance. Alternatively, other studies tried to correct astrocytic performance by boosting their normal functioning, for example by restoring glucose metabolism (Minhas et al., 2024) or overexpressing BDNF in astrocytes (de Pins et al., 2019). These studies also demonstrated positive effects on memory. We took a broader approach, and activated astrocytes via their Gq-coupled pathway, which increases intracellular Ca^2+^ and thus allows for a wide range of affects.

Normal memory has already been shown to improve by astrocytic Gq-coupled pathway activation in dorsal (Adamsky et al., 2018; Mederos et al., 2019; Refaeli et al., 2024) and ventral (Suthard et al., 2023) CA1. In one previous experiment using optogenetic stimulation of Gq-coupled pathway in astrocytes, no effect on hippocampal dependent memory was found in WT or 5XFAD mice, probably because the stimulation lasted only several minutes a day for 5 days (Gerasimov et al., 2023). Here we demonstrate memory enhancement in WTs in a new task, the RAWM, following hM3Dq activation in astrocytes, a finding similar to what we and others have shown before (Adamsky et al., 2018; Mederos et al., 2019; Refaeli et al., 2024; Suthard et al., 2023). Untreated 5XFAD mice perform worse than WTs, yet surprisingly, improve upon hM3Dq activation in astrocytes to the same levels as activated WTs.

One mechanistic explanation of the behavioral results is enhancement of neural activity and plasticity following astrocytic Gq-coupled pathway activation. Astrocyte effect on such processes has long been demonstrated (Perea et al., 2009). Specifically, the excitatory effect of astrocytic Gq-coupled pathway activation on neuronal activity and plasticity was first shown by us (Adamsky et al., 2018) and then by others (Gerasimov et al., 2023; Mederos et al., 2019; Van Den Herrewegen et al., 2021) in brain slices. Here, we show that LTP *in-vivo* in WTs is increased at least two-fold by hM3Dq stimulation. As has already been shown (Forest et al., 2021; Forner et al., 2021; MacPherson et al., 2017), 5XFAD mice have no LTP, yet upon astrocytic Gq-coupled pathway activation, normal LTP was observed.

Our finding of spontaneous neuronal hyper-activity in anesthetized 5XFAD mice and hypo-activity in awake behaving mice, is in line with a previous report showing that neural activity was also perturbed in a state-dependent manner in APP/PS1 mice (Zhou et al., 2022). The reported effects on neural activity in awake behaving 5XFAD mice may be due to the increased activation of plaque adjacent astrocytes, which were shown to be non-responsive to sensory stimulation near the plaque, and affect neuronal activity synchronization in the S1 cortex of APP/PS1 model mice (Lines et al., 2022). Additionally, in another AD model, PS2APP, astrocyte hypo-activity and neuronal LTP in slice were corrected by astrocytic STIM1 overexpression (Lia et al., 2023).

Another important function of astrocytes in AD is their capacity to internalize and degrade Aβ plaques (Giusti et al., 2024). Induction of non-specific phagocytosis of Aβ by astrocytes has been shown to reduce Aβ plaque burden in the hippocampus and PFC of 5XFAD mice and improve memory performance (Raha et al., 2021; Yang et al., 2024). Notably, our observations indicate that astrocytic phagocytosis of Aβ plaques is initiated as early as one day following activation of the Gq-pathway via CNO, consistent with previous reports of rapid plaque engulfment by astrocytes in the hippocampi of healthy mice (Prakash et al., 2021) and the cortex of 5XFAD mice (Iram et al., 2016). A previous study in APP/PS-1 mice showed that months-long Stat3 deletion in astrocytes improved memory and reduced Aβ plaque burden, but was caused by more Aβ internalization in microglial cells, rather than astrocytes (Reichenbach et al., 2019). Another study showed microglia internalize much more Aβ than astrocytes in healthy mice (Prakash et al., 2021). When we tested the phagocytic activity of microglia following Gq-pathway activation we found that on one hand, microglial Aβ internalization was much higher compared to that of astrocytes, but on the other, that it was not affected by astrocytic activation. Finally, we performed two experiments which bridge our results closer to possible clinical application. In the first, we evaluated the long-term effects of hM3Dq activation. Despite the fact the CNO is cleared from the body within hours, the beneficial effects on memory and physiology remain for two weeks in its absence. The effects, however, are not permanent, and disappear after 4 more weeks without CNO. A recent study identified a population of AD-associated astrocytes in 5XFAD mice that increase in quantity, at the expense of the normal population, as the disease progressed (Habib et al., 2020). In the future, it would be interesting to see whether activation via hM3Dq can bias astrocyte gene expression toward the healthy population, and whether this may have been the cause of the reported long-term effect without direct activation.

In the second experiment, we established that one year-long, continuous activation of the Gq pathway in astrocytes, from ages 7 to 19 months, is still effective, improves memory, reduces Aβ plaque burden, and elevates memory induced cFos. To the best of our knowledge, this is the only year-long continuous experiment studying the role of astrocytes in behavior and physiology, and the first in 5XFAD mice generally. It is important to show, as we did, that astrocytes can tolerate such a long activation.

As life expectancy rises, and global population ages, addressing the treatment of Alzheimer’s disease, the most common form of age-related dementias, presents a challenge of vital importance. Our work contrasts with most of the work in the field thus far, which conceptualized astrocytes as disease-promoting, and therefore attempted to block their activity in various ways ^e.g.^(Chen et al., 2025; Chicote-Gonzalez et al., 2025; Lee et al., 2023; Nakano-Kobayashi et al., 2023; Reichenbach et al., 2018; Zeng et al., 2023; Zhang et al., 2023). Instead, we report that activation of astrocytes via the Gq-pathway leads to a profound improvement in memory. This work also identifies Aβ plaque phagocytosis, accompanied by increase in neuronal activity and plasticity, suggesting potential mechanisms underlying astrocyte-driven improvement in memory in AD.

## AUTHOR CONTRIBUTION

T.K. performed behavior, histology, *in-vivo* electrophysiology and *in-vivo* 2-photon calcium imaging experiments; Y.W. contributed to behavioral experiments and histology. M.G. produced AAV vectors. A.D. contributed to 2-photon experiments; R.R. contributed to behavioral experiments; M.L. co-supervised electrophysiology experiments; I.G. conceived and supervised all aspects of the project, and wrote the manuscript with input from all other authors.

## ACKNOWLEDGMENTS

We thank the entire Goshen lab for their support. This project has received funding from the European Research Council (ERC) under the European Union’s Horizon 2020 research and innovation programme (grant agreement No 101087731 to I.G.). I.G. is also supported by the Israel Science Foundation (ISF grant No. 2060/23). M.L. is supported by the ISF (ISF grant No. 1331/23) and BSF. We thank Ami Citri, Neta Goshen and Nechama Novick for the critical reading of the manuscript. This work is dedicated to the memory of Mrs. Lily Safra, a great supporter of brain research.

## SUPPLEMENTARY FIGURES

**Figure S1:**
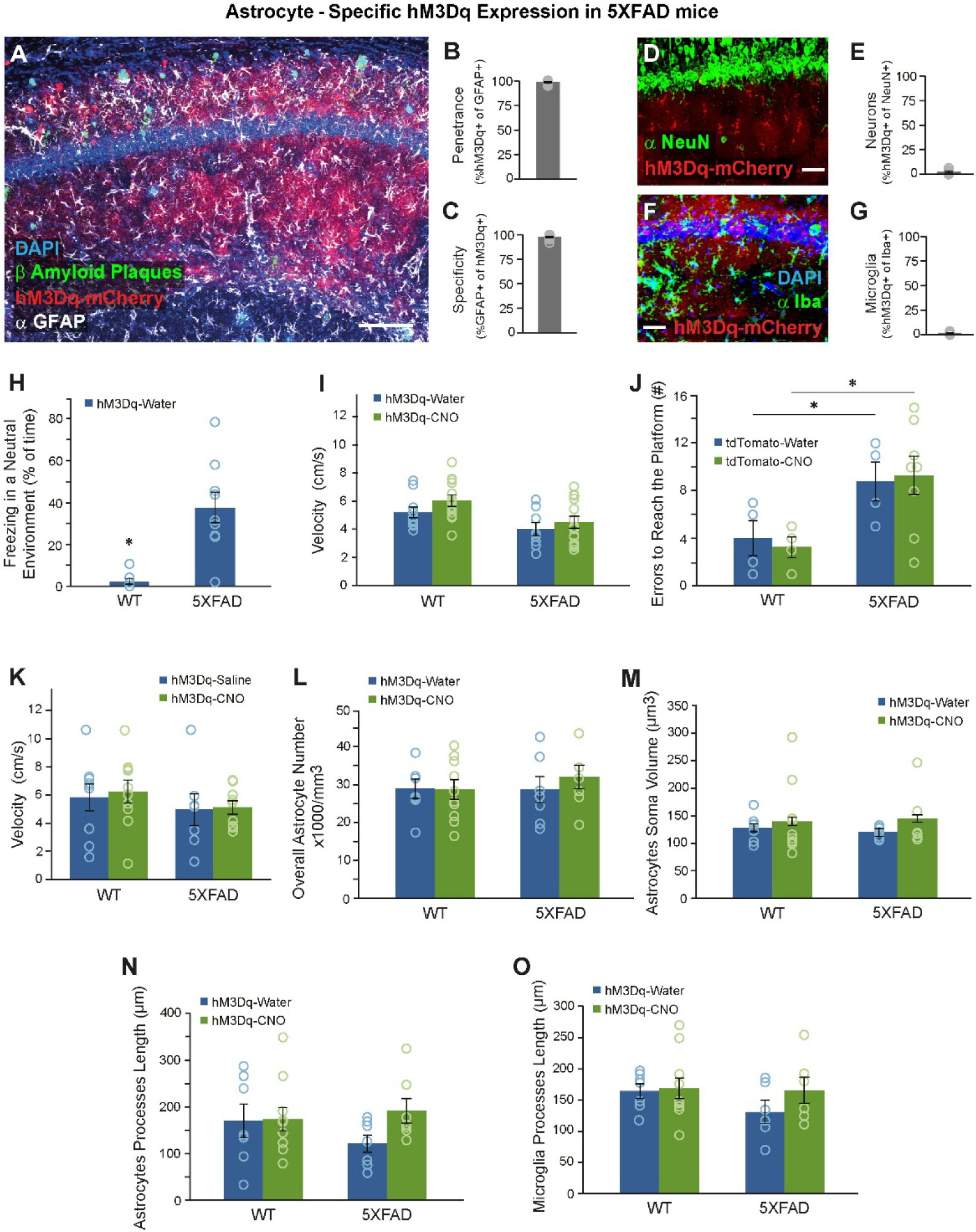
CNO application alone has no effect on memory or Aβ plaques in 5XFAD mice. (**A**) hM3Dq (red) was expressed in the astrocytes (stained with α-GFAP in white). Scale bar 100µm. (**B-C**) hM3Dq was expressed in >98% of CA1 astrocytes (1111/1127 cells from 4 mice), with >98% specificity (1111/1129 cells from 4 mice). No co-localization with the neuronal nuclear marker NeuN (**D-E**) or microglia marker Iba1 (**F-G**) was detected (scale bar 50µm). (**H**) 5XFAD mice (n=9) show freezing compared to WT (n=8), without conditioning (t(15)=4.442, p=0.000475). (**I**) Mice that drank either water (WT n=10; 5XFAD n=8) or CNO (WT n=13; 5XFAD n=13) swam at the same velocity. (**J**) In mice injected with a virus causing the expression of tdTomato, CNO (WT n=4; 5XFAD n=8) had no effect on the number of errors in the RAWM compared to water (WT n=4; 5XFAD n=4). Post hoc tests found effects of strain (5XFAD mice make more mistakes than WT) in Water (*= p<0.03) and CNO (*= p<0.015). (**K**) Mice that were injected for 4 days either with saline (WT n=9; 5XFAD n=7) or CNO (WT n=10; 5XFAD n=9) had the same velocity. (**L**) There was no effect on the overall number of astrocytes in hM3Dq-expressing mice that drank either water (WT n=7; 5XFAD n=7) or CNO (WT n=10; 5XFAD n=7). (**M**) There was no difference in astrocyte soma volume in hM3Dq-expressing mice that drank either water (WT n=7; 5XFAD n=7) or CNO (WT n=10; 5XFAD n=7). (**N**) There was no difference in astrocyte processes length in hM3Dq-expressing mice that drank either water (WT n=7; 5XFAD n=7) or CNO (WT n=10; 5XFAD n=7). (**O**) There was no difference in microglia processes length in hM3Dq-expressing mice that drank either water (WT n=7; 5XFAD n=6) or CNO (WT n=10; 5XFAD n=6). Data presented as mean ± standard error of the mean (SEM).

**Figure S2:**
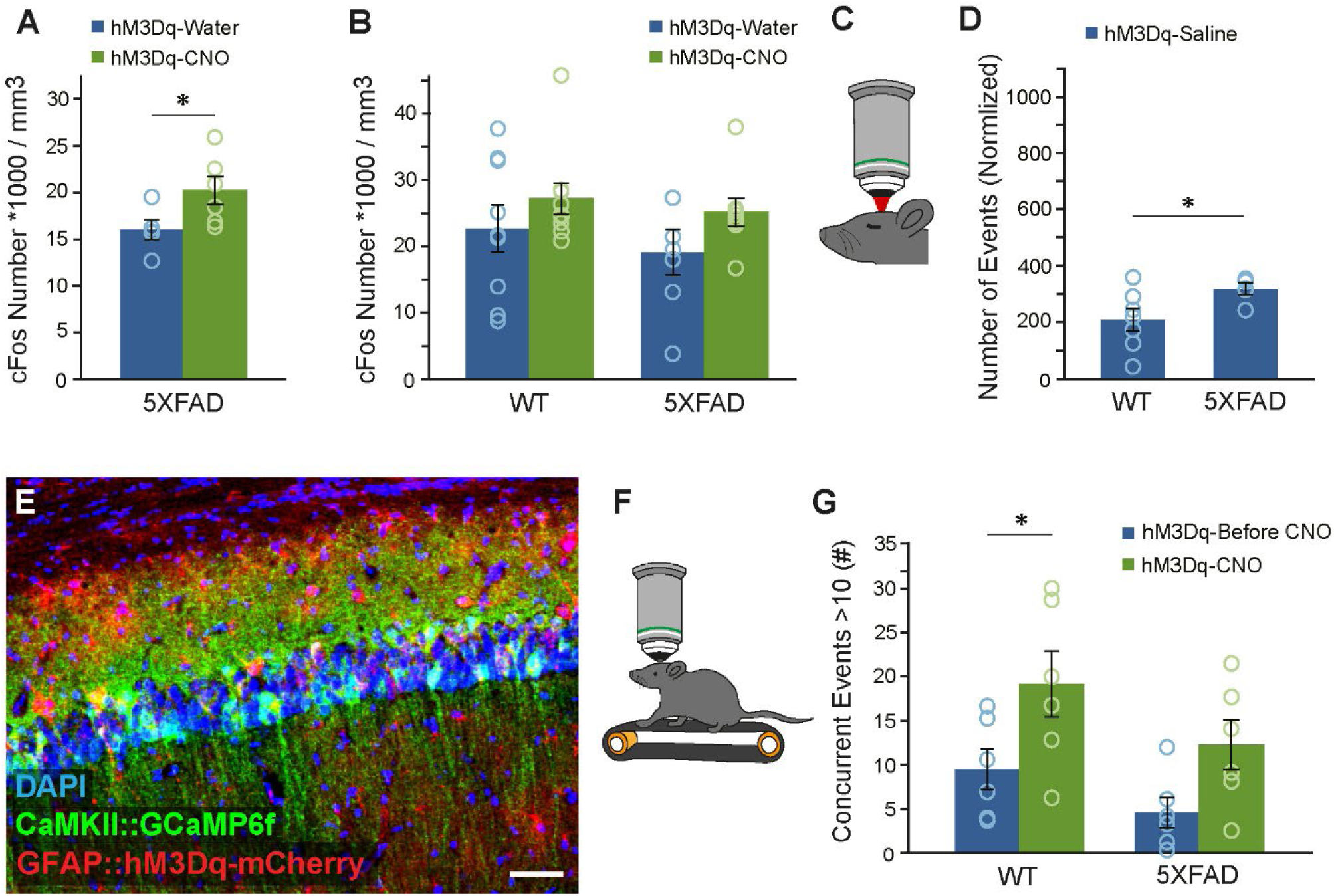
Neuronal activity is elevated in 5XFAD mice with astrocytic chemogenetic activation. (**A**) CNO application for 2 weeks significantly elevated cFos levels in 5XFAD female mice (Water n=5; CNO n=6)(*= p< 0.03). (**B**) CNO injections for 4 days significantly elevated cFos levels overall (WT-saline n=9; WT-CNO n=9; 5XFAD-saline n=6; 5XFAD-CNO n=8). (**C**) Ca^2+^ events in neurons were measured *in-vivo*, in anesthetized mice. (**D**) Anesthetized 5XFAD mice (n=5) showed increased number of events compared to anesthetized WTs (n=7)(*= p< 0.03). (**E**) Co-expression of hM3Dq expression (red) in astrocytes, and GCaMP6F in neurons (green). Nuclei are in blue. Scale bar 50µm. (**F**) Ca^2+^ events in neurons were measured, *in-vivo*, in awake mice. (**G**) CNO application increased the number of concurrent events overall (p=0.002). Post hoc analysis showed that CNO significantly elevated the number of concurrent events in WT mice (n=6) (*= p< 0.01), but not in 5XFAD mice (n=6). Data presented as mean ± standard error of the mean (SEM).

**Figure S3:**
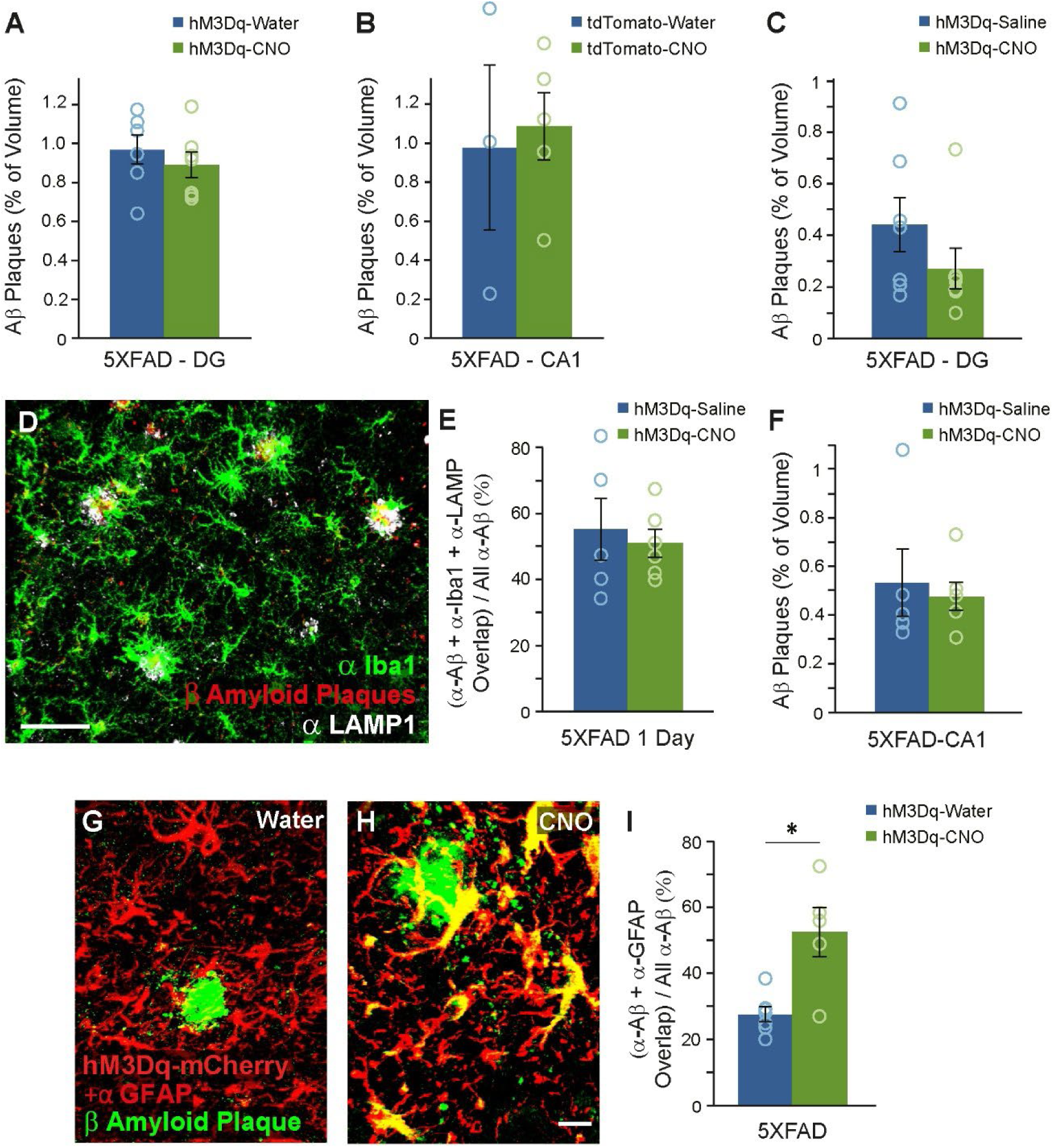
No effect on microglia endocytosis of Aβ plaques by chemogenetic astrocytic Gq-pathway activation in an Alzheimer’s disease model. (**A**) CNO-treated 5XFAD mice (n=7) expressing hM3Dq in their CA1 astrocytes showed no difference in their Aβ plaque in the DG compared to 5XFAD mice that received water (n=6). (**B**) CNO (n=5) or water (n=3) treated 5XFAD mice with a control tdTomato only virus showed no difference in the Aβ plaques in their CA1. (**C**) CNO-injected 5XFAD mice (n=7) expressing hM3Dq in their CA1 astrocytes showed no difference in Aβ plaques in their DG compared to 5XFAD mice that were injected with saline (n=7). (**D**) Staining for microglia (α-Iba1; green), Aβ plaques (red) and lysosomes (α-LAMP1; white). Scale bar 30µm. (**E**) One day following injection, 5XFAD-CNO (n=6) showed no change in overlap of microglia, Aβ plaques and lysosomes compared to 5XFAD-saline (n=5). (**F**) One day following injection, no significant difference in Aβ plaque volume was found between 5XFAD-CNO (n=6) and 5XFAD-saline (n=5). Data presented as mean ± standard error of the mean (SEM). (**G-H**) Expression of hM3Dq + GFAP staining (red) and staining for Aβ plaques (green). Scale bar 10µm. (**I**) After 2 weeks of CNO in the drinking water, 5XFAD-CNO (n=5) showed 90% more overlap of astrocytes and Aβ plaques compared to 5XFAD-saline (n=7)(*= p< 0.002). Data presented as mean ± standard error of the mean (SEM).

**Figure S4:**
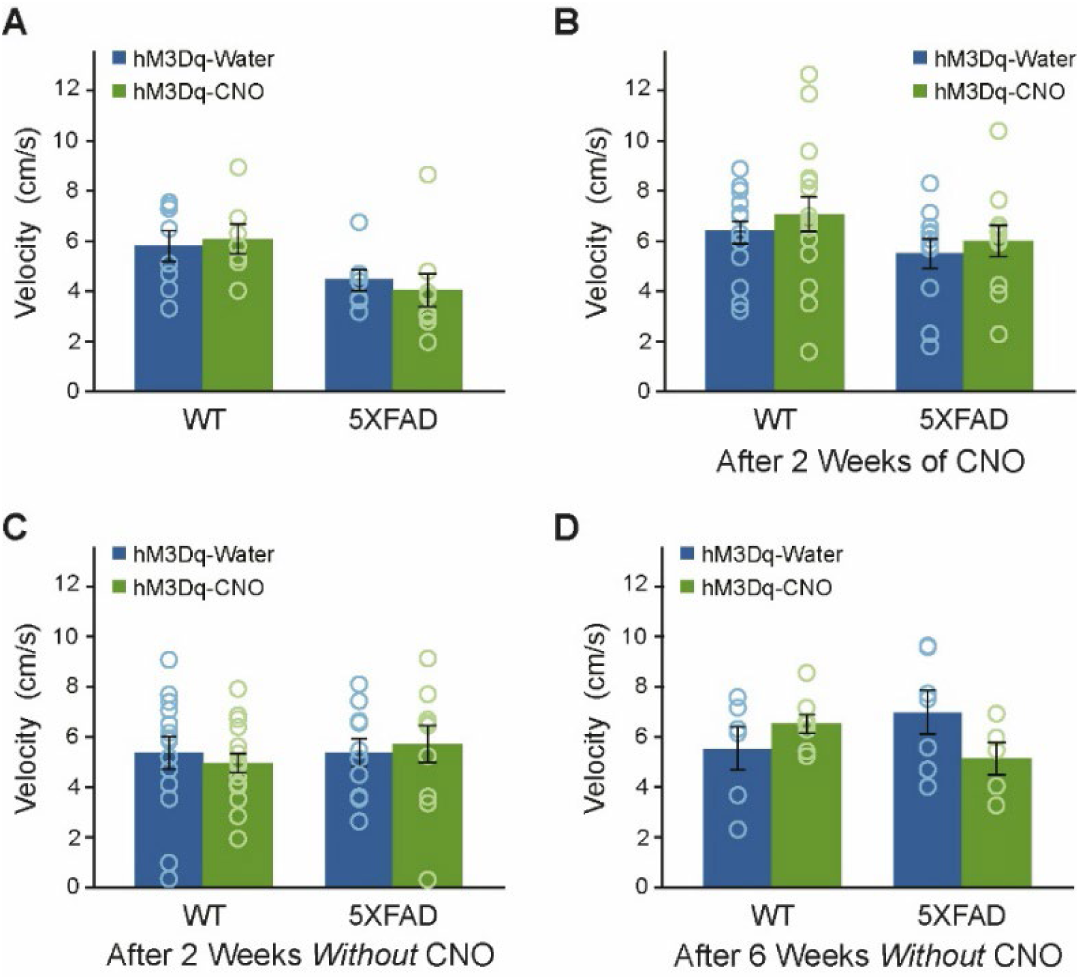
Mice tested in the clinically relevant schedules showed no effect on their velocity. (**A**) Mice that drank either Water (WT n=8; 5XFAD n=7) or CNO (WT n=7; 5XFAD n=8) for a year had the same velocity. (**B**) Mice that drank either water (WT n=14; 5XFAD n=11) or CNO (WT n=16; 5XFAD n=11) for two weeks swam at the same velocity. (**C**) After two weeks *without* CNO, the same mice still performed at the same velocity. (**D**) After six weeks *without* CNO, mice that drank either Water (WT n=6; 5XFAD n=7) or CNO (WT n=8; 5XFAD n=5) at the beginning of the experiment showed the same velocity. Data presented as mean ± standard error of the mean (SEM).

## METHODS

### Mice

5XFAD (Tg(APPSwFlLon,PSEN1*M146L*L286V)6799Vas/Mmjax) and their WT littermates, were bred in our SPF animal facility. They were bred with a heterozygous 5XFAD male and a WT female, as recommended (Sasmita et al., 2025), and the littermates were genotyped.

All experimental protocols were approved by the Hebrew University Animal Care and Use Committee.

### Stereotactic Injections

Mice were anesthetized with isoflurane, and their heads placed in a stereotactic apparatus (Kopf Instruments, USA). The skull was exposed and a small craniotomy was performed. To cover the entire dorsal CA1, mice were bilaterally microinjected in two sites per hemisphere using the following coordinates: Site 1: anterior–posterior (AP) −1.5mm from bregma, medial–lateral (ML) ±1mm, dorsal–ventral (DV) −1.55mm; Site 2: AP −2.5mm, ML ±2mm, DV −1.55mm. Microinjections were performed using a 10µL syringe and a 34-gauge metal needle (WPI, Sarasota, USA). The injection volume and flow rate (0.1ml/min) were controlled by an injection pump (WPI). Following each injection, the needle was left in place for 10 additional minutes to allow for diffusion of the viral vector away from the needle track and was then slowly withdrawn. The incision was closed using sewing and Vetbond tissue adhesive. For postoperative care, mice were subcutaneously injected with Carprofen (5mg/kg).

For 2 photon imagining mice were unilaterally microinjected 400 nl viral vector using the following dorsal CA1 coordinates: anteroposterior, −1.85 mm; mediolateral, +1.4 mm; and dorsoventral, −1.45 mm from bregma. All microinjections were performed using a 10 µl syringe and a 34-gauge metal needle (WPI). The injection volume and flow rate (0.1 µl min−1) were controlled by an injection pump (WPI). After each injection, the needle was left in place for an additional 10 min to allow for diffusion of the viral vector away from the needle track, and was then slowly withdrawn. The craniotomy was sealed with bone wax (Surgical Specialties), and the exposed skull was covered with transparent super-bond (Sun Medical) for cementing an omega-shaped head bar (custom design, 3D printed) anteriorly to the craniotomy site. For postoperative care, the mice were subcutaneously injected with Carprofen (5mg/kg).

After at least one week of rest, mice were re-anaesthetized with isoflurane in the stereotactic apparatus, and a biopsy punch (Kai Medical) was used to cut a ∼2.5 mm diameter craniotomy over the injection site. Aspiration was used to remove the cortical tissue and top most fibres above the right dorsal CA1, and a glass cannula (2.4 mm diameter, 2.5 mm length, no. 0 cover slip bottom; self-fabricated) was inserted into the craniotomy. The skull was covered with opaque super-bond (Sun Medical) for cementing the cannula. An additional layer of dental acrylic was placed to minimize potential physical damage.

### Viral Vectors

The following dilutions and volumes of vectors were used: AAV1-gfaABC1D-hM3Dq-mCherry (EVCF, titer 1.5E13, 700nl/site), AAV5.CaMK2.GCaMP6f.WPRE.SV40 (Addgene titer 3.1E13) together with AAV1-gfaABC1D-hM3Dq-mCherry (EVCF, titer 1.5E13) in a ratio of 1:1 (500nl), AAV1-gfaABC1D-tdTomato (EVCF, titer 5.6E13, 600nl/site)

### CNO Administration

*CNO in drinking water:* For chronic activation of astrocytes, 22mg CNO (Tocris #4936) were dissolved in 1ml of DMSO and added along with 10g sucrose into 1L of water. Drinking bottles were replaced every 24hr and protected from light.

*IP injections*: CNO was dissolved in DMSO and diluted in 0.9% saline to yield a final DMSO concentration of 0.5%. Saline solution for control injections also consisted of 0.5% DMSO. 3mg/kg CNO was intraperitoneally injected 30min before the behavioral assays.

### Radial arm water maze (RAWM)

The apparatus consists of a water pool (85cm in diameter) with six 14cm wide arms radiating from the central circular area. The escape platform (11X11) was placed in a different arm during each of 5 consecutive days of acquisition (Licht et al., 2020), with 5 repetitions per day. At the beginning of each acquisition trial, the animal was placed in a different arm (excluding the escape platform) and allowed 1 min to swim into any of the arms until finding the platform. If during this time the platform was not found, the mouse was gently placed on the platform and allowed to stay there for 10sec. The fifth trial on the fifth day was the test trial, in which the number of errors to reach the platform were calculated. Each error was defined as follows: (1) swimming into an arm that does not contain the platform that day (1 error for every wrong arm entrance), (2) entering the goal arm without boarding the platform, or (3) spending ≥20 s continuously in the central zone without any arm selection.

All trials were recorded and analyzed using EthoVision XT software (Noldus). After swimming, mice were dried using a heating lamp.

### Immunohistochemistry

After the last behavioral experiment, mice were transcardially perfused with cold PBS followed by 4% paraformaldehyde (PFA) in PBS. The brains were extracted, postfixed overnight in 4% PFA at 4°C and cryoprotected in 30% sucrose in PBS. Brains were sectioned to a thickness of 40 μm using a sliding freezing microtome (Leica SM 2010R) and preserved in a cryoprotectant solution (25% glycerol and 30% ethylene glycol in PBS). Free-floating sections were washed in PBS and incubated for 1 h in blocking solution (1% of bovine serum albumin, BSA, and 0.3% Triton X-100 in PBS) at room temperature.

For cFos staining, the relevant brain slices were incubated for 7 days at 4°C with rabbit anti-cFos primary antibody (Synaptic system, #226003). For all the other staining, slices were incubated overnight at 4°C (Rabbit anti IBA-1, Wako# 019-19741; Rabbit anti Aβ, Merck #AB5074P; Chicken anti GFAP, Merck #AB5541, Mouse anti Aβ, BioLegend#383002; Rat anti Lamp-1, Abcam #25245). Sections were then washed with PBS and incubated for 2 hr at room temperature with secondary antibody (1:500 donkey anti rabbit, donkey anti mouse, donkey anti rat, donkey anti chicken, Jackson laboratory) in 1% BSA in PBS. Finally, sections were washed in PBS, incubated with 4,6-diamidino-2-phenylindole (DAPI; Sigma D9542, 1μg ml^-1^), and mounted on slides with Mounting Medium (Dako, #S3025).

### Confocal Imaging and Quantification

Confocal fluorescence images (40um) were acquired using an Olympus scanning laser microscope Fluoview FV1000 using 4X and 10X air objectives, or 20X and 40X oil immersion objectives. Images were analyzed using the IMARIS 10 software (Bitplane, UK).

Using the ‘surfaces” feature in the IMARIS software, for each slice, all Aβ plaques were marked and their volume was calculated by the software. The volume of the plaques was divided by the measured volume of the CA1 in the slice.

For astrocytes and microglia soma body and processes measurements, we used the ‘Filaments’ feature in the IMARIS software.

For measurements of Lysosomes and Aβ inside astrocytes or microglia, or the measurement of Aβ inside astrocytes a colocalization channel of the 2 or 3 different staining colors was performed for each slice using the IMARIS software. Then, using the ‘surfaces” feature in the IMARIS software, co-localized stainings were marked and their volume was calculated by the software. The volume of the Lysosomes + astrocytes/microglia + Aβ, or astrocytes + Aβ inside astrocytes was divided by the measured Aβ plaques volume of that slice.

### LTP in anesthetized mice

Mice were anesthetized with isoflurane and 2 electrodes glued together were lowered into their brain. A tungsten microelectrode (1MΩ, ∼125µm, A-M systems) was used to record local field potentials in the stratum radiatum (AP: −1.5mm; ML: 1.1mm; DV: −1.5mm). A sereotrode (1MΩ, 2-3 µm, WPI) was placed (AP: −1.5mm; ML: 1.8mm; DV: −1.7mm) to stimulate the Schaffer collaterals. saline or CNO were injected 30min before the HFS protocol started. After 15min of baseline recording (one 0.1mA stimulus every 20s), we waited 15min, and then each animal was presented with a 10 minutes HFS protocol consisting of five trains (200Hz, 100ms; 1 per second), presented six times at 1min intervals. Evolution of fEPSPs in response to the HFS was recorded after 60 minutes for 15min (one 0.1 mA stimulus every 20s). Recordings was carried out using a Multiclamp 700B amplifier (Molecular Devices). Signals were low-pass filtered at 5kHz, digitized and sampled through an AD converter (Molecular Devices) at 10kHz and stored for offline analysis using Matlab (Mathworks). CA1 responses to Schaffer collateral stimulation were quantified by calculating the slope of the each fEPSPs relative to the mean baseline levels.

### 2-Photon Microscope

Imaging was performed using the Neurolabware 2-photon laser scanning microscope (Los Angeles, CA, USA). Excitation light from a Ti:sapphire laser (Chameleon Discovery TPC, Coherent) operated at 920 nm scanned the sample using a 6215 galvometer and a CRS8 resonant mirror (Cambridge Technology). Emitted fluorescence light was detected by GaAsP photo-multiplier tubes (Hamamatsu, H10770-40) after bandpass filtering (Semrock). XYZ motion control was obtained using motorized linear stages, enabled via an electronic rotary encoder (KnobbyII). A molding clay ring was mounted between the cannula and the objective in order to maintain the water reservoir and block external light. The Scanbox software, run on MATLAB, was used for microscope control and image acquisition. All images were acquired using a water immersion 16X objective (Nikon, 0.8 NA) with magnification of 2.8 to obtain 601×418µm fields of view. The sampling rate was 15.49 frames per second.

To ensure that we image the same field of view over sessions, for the experiments measuring neuronal activity before and during CNO in awake and anaesthetized mice (1% Isoflorane) mice were not removed from the 2-photon setup during the whole imaging session.

### Processing Calcium Imaging Data

Suite2p was used to process two-photon calcium imaging data (Pachitariu et al., 2017). Movement correction was done using non-rigid motion-correction with 1000 frames used to compute the reference image. Cell detection was performed using the default suite2p classifier. Neuropil correction and classification was done using the default suite2P parameters (https://suite2p.readthedocs.io/en/latest/settings.html).

*ROI Detection:* The movies of before and during CNO in awake and anaesthetized mice were analyzed together and only ROIs that were visible in both imaging sessions were included in the analysis. We used the Suite2P software to detect ROIs based on their morphology and activity pattern, and than manually checked each ROI to ensure that they were indeed neurons.

*Signal Extraction:* The mean fluorescence intensity time series were extracted from the ROIs using Suite2P and analyzed with MATLAB. The first five samples were removed from the analysis. To obtain Δ*F*/*F* time series for each ROI, we adapted previously published methods for the neuronal signal (Dombeck et al., 2010). We defined the baseline F as the eighth percentile fluorescence value within a 15s interval around each sample point. We then subtracted the baseline from each sample, and divided the result by the baseline F. Noisy ROIs, in which there were no apparent calcium transients, were removed from the analysis.

*Detection of calcium events*: We used previously published methods to identify significant calcium events (Dombeck et al., 2010). The Δ*F*/*F* traces were expressed as the standard deviations from the baseline fluorescence. Event onset was defined as fluorescence >2σ and the offset as fluorescence <0.5σ. We calculated the ratio between negative and positive events as a function of length. Events whose duration exceeded the minimal event length at which the ratio <0.01 were considered significant. The procedure was performed in 3 iterations. Initially the σ was defined for the entire timeseries, and in each of the following iterations the previously detected significant transients were excluded when calculating σ.

### Quantification and statistical analysis

All data were processed and analyzed using Excel, SPSS and MATLAB. For comparisons we preformed One-way or repeated measures ANOVA with LSD post hoc, and one or two tailed Student’s t test as described in the corresponding figure legend. All statistical details of experiments can be found in the results, figures and figure legends.

The results in Figures were presented as group mean and standard error of the mean (SEM). Statistical significance is indicated with asterisks as; ∗ p < 0.05.

## Notes

### Competing Interest Statement

The authors have declared no competing interest.

